# Membrane Biofouling and Plasmid Conformation Shape the Retention of Antibiotic Resistance Gene-Carrying Plasmids during Ultrafiltration

**DOI:** 10.64898/2026.11.17.767232

**Authors:** Wenxuan Chen, Fátima Rojas Serrano, René Kallies, Susann Müller, Niculina Musat, Florian Schattenberg, Matthias Schmidt, Hendrik Seifert, Gangan Wang, Anja Worrich

## Abstract

Antimicrobial resistance poses a growing threat to human and environmental health, and wastewater treatment plants are important pathways for the release of antibiotic resistance genes (ARGs). Although ultrafiltration (UF) membranes effectively retain antibiotic-resistant bacteria, their capacity to remove extracellular ARGs remains less understood. Extracellular plasmids are concerning because they persist following cell lysis and may contribute to horizontal gene transfer. Unlike intact bacterial cells, these flexible DNA molecules may pass through membrane barriers. Here, we investigated the removal of extracellular plasmids carrying ARGs using a 100 kDa molecular weight cut-off (MWCO) UF membrane, focusing on biofouling, plasmid size, and conformation. Initial filtration experiments achieved removal efficiency (*RE*) above 99.9% for antibiotic-resistant bacteria and plasmids, and produced a pronounced biofouling layer. For the pre-fouled membrane, mass balances showed increased adsorption and permeation of plasmids, with *RE* declining from approximately 99% to below 95% over consecutive runs. Microscopy indicated plasmid deposition in biofouled regions. Additionally, plasmid size showed no consistent relationship with *RE*, while conformation had a pronounced effect, with supercoiled plasmids being retained more efficiently (98.8–99.6%) than their linear counterparts (82.4–92.4%). These findings demonstrate that plasmid retention by UF is influenced by both membrane conditions and DNA conformation, which nominal membrane pore size alone cannot capture. The high *RE* highlights the potential of UF as a tertiary treatment, while emphasizing the need to account for biofouling and DNA structural diversity when evaluating strategies to reduce extracellular ARG discharge from wastewater treatment plants.

## 1. Introduction

The widespread use of antibiotics in human and veterinary medicine has accelerated the emergence and dissemination of antibiotic resistance, posing a major threat to global health (Murray et al., 2022; O’Neill, 2016). Wastewater treatment plants (WWTPs) receive antibiotics, antibiotic-resistant bacteria (ARB), and antibiotic resistance genes (ARGs) from households, hospitals, and industrial sources and represent important pathways for antimicrobial resistance dissemination (Drane et al., 2024). Although conventional activated sludge systems efficiently remove organic matter and nutrients, ARB and ARGs are frequently detected in treated effluents and receiving water bodies (Czekalski et al., 2014; Rodriguez-Mozaz et al., 2015; Su et al., 2024). This concern is reflected in the revised European Urban Wastewater Treatment Directive, which introduces monitoring requirements for antimicrobial resistance in large WWTPs. However, monitoring alone cannot prevent the release of resistance determinants, highlighting the need for effective treatment strategies.

Membrane-based treatment technologies offer promising opportunities to reduce the discharge of ARB and ARGs (Mardalisa et al., 2023). Membrane bioreactors (MBRs) generally achieve higher removal efficiencies (*RE*s) than conventional treatment (Cheng and Hong, 2017; Le et al., 2018), but replacing existing WWTP infrastructure with MBRs is often economically impractical. Ultrafiltration (UF) as a tertiary treatment step could therefore provide a more feasible alternative, particularly because membrane filtration efficiently retains intact bacterial cells (Schwermer et al., 2018). However, ARGs are not exclusively associated with intact bacteria. Bacterial cell lysis during wastewater treatment releases DNA, including plasmids carrying ARGs, into the aqueous phase (Calderón-Franco et al., 2021). Such release may also occur during oxidative or disinfection treatments, including ozonation (Zhuang et al., 2015). Unlike bacterial cells, extracellular plasmids are much smaller and may exhibit fundamentally different filtration behavior (Slipko et al., 2019). Moreover, extracellular plasmid DNA can remain biologically relevant by contributing to horizontal gene transfer (HGT) in receiving environments (Calderón-Franco et al., 2021). Efficient bacterial removal therefore does not necessarily imply equally efficient removal of extracellular resistance determinants.

Plasmids are important carriers of ARGs and contribute to their dissemination across bacterial communities (Rozwandowicz et al., 2018; Zhang et al., 2023). Although membrane processes are widely used for plasmid purification in biotechnology (Affandy et al., 2013; Arkhangelsky et al., 2011), relatively little is known about plasmid retention during wastewater treatment. One potentially important factor is membrane fouling, which can alter membrane permeability, surface interactions, and solute transport (Hajibabania et al., 2011). Studies using MBR systems have reported substantial, but sometimes inconsistent, effects of fouling on extracellular ARG removal (Breazeal et al., 2013; Latulippe and Zydney, 2008). However, tertiary UF systems differ from MBRs in their hydrodynamic conditions, fouling dynamics, and medium composition, making it difficult to directly transfer these findings. Consequently, the influence of biofouling on extracellular plasmid retention during tertiary UF remains insufficiently understood.

Beyond membrane conditions, the physicochemical properties of plasmids may strongly influence their retention. Most studies have focused on membrane pore size or molecular weight cut-off (MWCO) (Breazeal et al., 2013; Krzeminski et al., 2020; Slipko et al., 2019), although DNA molecules behave differently from rigid particles because of their flexibility and ability to deform under flow (Morão et al., 2011). In particular, plasmid conformation may influence transport through membrane pores: supercoiled plasmids adopt relatively compact, branched structures, whereas linear DNA can elongate more readily under hydrodynamic stress (Li et al., 2016). Plasmid size, i.e., the length in base pairs (bp) may further affect these processes, although it alone may not predict retention efficiency, as the effective molecular size also depends on DNA conformation and base content. Importantly, extracellular DNA in treated wastewater is unlikely to occur exclusively in an intact supercoiled state. Cell lysis and treatment-related stresses can generate open circular, linearized, or fragmented DNA (Slipko et al., 2019). While these forms may differ in their capacity for HGT, linear DNA can still contribute to genetic exchange through recombination-based mechanisms. Understanding how plasmid size and conformation affect membrane retention is therefore important for evaluating the potential of UF to limit extracellular ARG dissemination.

In this study, we investigated the potential of a 100 kDa MWCO UF membrane for retaining extracellular plasmids carrying ARGs under conditions relevant to tertiary wastewater treatment. We focused on two complementary determinants of plasmid retention: membrane biofouling and plasmid-intrinsic properties. Using a laboratory-scale cross-flow filtration system, we compared plasmid removal by initially non-fouled and pre-fouled membranes and examined the effects of plasmid size and conformation (supercoiled versus linear) under initially non-fouled conditions. By combining *RE* measurements, plasmid mass balance analyses, and membrane imaging, we aimed to characterize how these factors influence plasmid rejection, membrane-associated retention, and permeation, thereby improving our understanding of extracellular ARG removal by tertiary UF.

## 2. Materials and Methods

### 2.1. Study design

The study was divided into two complementary stages. The first investigated the influence of membrane biofouling on plasmid retention, whereas the second examined the influence of plasmid-intrinsic properties, namely size and conformation, on UF performance.

Part I included two batches of experiments, one with a new membrane cassette and a second with a membrane cassette that had previously been used to filter a bacterial solution. Each batch consisted of seven runs performed on consecutive days, with the aim of comparing how the membrane *RE* evolves over time under different fouling conditions. Part II included six experiments, as three different sized plasmids were tested in two conformations (supercoiled and linear). A new membrane cassette was used for each plasmid size–conformation pair.

### 2.2. Preparation of the synthetic effluent medium and model plasmids

A synthetic effluent medium was inoculated with the model plasmids. The medium was prepared as a dilution of an artificial wastewater medium described by Riedl et al. (2017), with the exact medium composition described in Supporting Information 1 (SI1).

All the plasmids used in this study harbor resistance genes to some of the most common antibiotics found in wastewater (Rodriguez-Mozaz et al., 2015). For fouling experiments, plasmid R388 (33.9 kbp), conferring resistance to sulfonamides (Garcillán-Barcia et al., 2011) and trimethoprim, was used as a model plasmid to evaluate the effect of membrane fouling on plasmid removal. This plasmid is broad-host-range and belongs to the group of conjugative plasmids studied in Enterobacteriaceae (Avila and de la Cruz, 1988), widely found in wastewater (Cangola et al., 2025; Silva et al., 2007). To test the effects of plasmid size and conformation, three different plasmids were used: pDsRed2 (https://www.snapgene.com/plasmids), pUX-BF13 (Bao et al., 1991) conferring resistance to ampicillin, and RSF1010 (Garcillán-Barcia et al., 2011) conferring resistance to sulfonamides. These plasmids ranged from 3 to 12 kbp, and different conformations were obtained by linearizing plasmids by cutting them with restriction enzymes (Table 1). Here, plasmid size refers to the length in bp, whereas conformation refers to the topology of the plasmid. The extraction and linearization procedures of the extracted plasmids can be found in SI2.

**Table 1.** The characteristics of the plasmids together with the respective strains.

| **Plasmid (conformation)** | **Harboring strain** | **Factor studied** | **Size (bp)** | **Target ARG** | **Concentration in feed (ng/L)** |
|---|---|---|---|---|---|
| R388 (supercoiled) | E. coli K 12 J53 | Fouling (new/fouled membrane) | 33926 | *sul1* | 6188 |
| pDsRed2 (supercoiled) | E. coli DH5α | Plasmid size and conformation | 3311 | *blaTEM* | 10993 |
| pDsRed2 (linear) | E. coli DH5α | Plasmid size and conformation | 3311 | *blaTEM* | 12589 |
| RSF1010 (supercoiled) | E. coli K12C600 | Plasmid size and conformation | 8684 | *sul2* | 5233 |
| RSF1010 (linear) | E. coli K12C600 | Plasmid size and conformation | 8684 | *sul2* | 6279 |
| pUX-BF13 (supercoiled) | E. coli SM10 λpir | Plasmid size and conformation | 12256 | *blaTEM* | 9711 |
| pUX-BF13 (linear) | E. coli SM10 λpir | Plasmid size and conformation | 12256 | *blaTEM* | 13950 |

A volume of 4.3 L autoclaved synthetic effluent medium was used to dissolve the respective plasmid types. Subsequent mixing (1000 rpm) for 20 min using magnetic stirrer bars placed in the bottles to avoid plasmid deposition and guarantee a homogeneous mixture. The plasmid concentrations in the feed varied for the different experiments due to variable extraction yields but they always ranged between 5000 and 14000 ng/L (Table 1), which are extracellular ARG concentrations typically found in secondary effluents (Calderón-Franco et al., 2021; Li et al., 2019; Zhang et al., 2018). The plasmid concentration in the feed was always constant for the consecutive runs in the same batch of experiments.

The European Urban Wastewater Treatment Directive establishes the Chemical Oxygen Demand (COD) of 125 mg/L O_2_ as the maximum limit for wastewater discharge of secondary effluents. To guarantee a synthetic effluent medium quality comparable to that of secondary effluents, the COD was measured before and after the plasmid addition (SI1).

### 2.3. Experimental facility and membrane filtration experiments

The experimental facility (SI3) included a membrane cassette, Vivaflow 200 Laboratory Cross Flow (VF20P4, Sartorius, Germany), 100 kDa MWCO, connected to an autoclaved 5 L feed bottle by a peristaltic pump (Masterflex® L/S®, Cole-Parmer, USA). The feed was continuously mixed at 800 rpm using magnetic stirrer to avoid plasmid deposition during the experiments. The retentate was recirculated to the feed bottle, while the permeate was collected in a separate bottle. The feed flow rate was constant for all the experiments and took values of approximately 300 mL/min. The permeate weight was measured with a balance (MS6002TS/00, Mettler-Toledo International Inc, Switzerland) connected to a computer, which allowed the registration of the permeate flow rate with BalanceLink software (Mettler Toledo, Switzerland). Pressure gauges in the feed and reject streams allowed measuring the pressure, whereas the temperature was maintained at 25°C as the lab-scale plant was located inside an incubator (Unihood 650, UniEquip Laborgerätebau und Vertriebs GmbH, Germany).

Before use, every new membrane was subject to compaction, a common practice to unpack the pores and remove preservatives (Arenillas et al., 2017). Each membrane was soaked in autoclaved Milli-Q® water overnight, and the day after, a blank filtration with 30 L of autoclaved Milli-Q® water was performed before the plasmid experiments.

Every filtration run finished when the permeate volume reached 3.8 L. Triplicate samples of feed and permeate were collected in 1.5 mL low-binding tubes at the start and at the end of each run to calculate the membrane *RE* with Eq.1 (section 2.7). The samples were taken by means of Discofix® C three way-valves with sterile syringes. Afterwards, they were stored at 4°C before being analyzed. In addition, the membrane was subjected to blank filtrations with autoclaved Milli-Q® water before and after synthetic effluent medium filtration. Permeate samples were collected after the blank filtration preceding plasmid filtration and analyzed accordingly to verify there was no bias between runs.

### 2.4. ARGs quantification

The plasmids tested contained *sul1*, *sul2* and *blaTEM* resistance genes, as indicated in Table 1. The copy number of the respected amplicons was quantified by SYBRGreen quantitative polymerase chain reaction (qPCR) with the feed and permeate samples collected during the UF runs. All the reactions were performed in a CFX Opus 96 Real-Time PCR System cycler (Bio-Rad Laboratories, Inc, USA) with a 12.5 µL reaction mixture containing 6.25 μl SsoAdvanced Universal SYBR Green Supermix (Bio-Rad Laboratories, Inc, USA), 0.25 μl (2.5 pmol, 200 nM) forward primer, 0.25 μl (2.5 pmol, 200 nM) reverse primer, 3.25 μl PCR water, and 2.5 μL of the respective sample or standard. The qPCR protocol comprised a denaturation step at 98 °C for 2 min, followed by 40 cycles of amplification. The different times and temperature values applied for the amplification of each target gene together with the primers used can be found, respectively, in Tables S3 and S4.

ARG copy numbers were quantified using assay-specific calibration curves generated from five independent qPCR runs, each including standards measured in triplicate. Amplification efficiencies ranged from 90% to 110%. Calibration parameters, limits of detection (LoD), and limits of quantification (LoQ) are provided in the Table S5 and S6.

### 2.5. Establishment of a fouling layer on the membrane

To evaluate the effects of fouling on plasmid retention, a new membrane was previously fouled by filtering fresh bacterial solutions on 3 consecutive days per week over a 4 weeks period. *Pseudomonas putida* KT2440, a well-characterized model strain of a genus commonly found in WWTPs (Amiri et al., 2022; Lyu et al., 2024), was selected as model bacterium (SI5). The feed had a bacterial cell concentration of 2×10^7^ cells/mL, which lies within the range reported for WWTP effluents (10^6^−10^8^ cells/mL) (Foladori et al., 2010; Ma et al., 2013; Manti et al., 2008).

The feed and permeate samples were collected at the beginning and end of each UF run and analyzed by flow cytometry (SI6). Flow cytometry is an often-used technology to count bacterial concentrations in water environments (Vives-Rego et al., 2000). First, samples were nucleic acid stained with SYTO®9 (Lot. 2088729; ThermoFisher Scientific, Eugene, OR, USA) as in Li et al. (2022). Cell counting was done as in Liu et al. (2019). These analyses were conducted to verify the retention of bacteria responsible for biofilm formation on the UF surface to confirm the reliable functioning of the filtration system during biofouling. After 12 UF runs, the membrane cassette was opened to extract the membrane sheet for a membrane autopsy to confirm the development of biofouling (section 2.6). Since a membrane autopsy is a destructive sampling, an independent pre-fouling experiment was necessary to obtain the pre-fouled membrane to be used in the subsequent plasmid experiments (section 2.2). An analogous experiment was performed, but the bacterial concentration of *P. putida* KT2440 was increased to 3×10^7^ cells/mL, with 4 consecutive runs per week for 2 weeks. In this way, the number of bacteria filtered was the same, but the duration was reduced by half. The device was then disassembled and the membrane cassette stored at 4°C until the start of the plasmid experiments. More details about respective pre-experiments can be found in the SI5 and Fig. S5.

### 2.6. Fouling visualization after membrane autopsy

After the UF experiments had been completed, the respective cassettes were opened and the membrane sheets were extracted for microscopy analyses. Helium Ion Microscopy (HIM) or field-emission scanning electron microscopy (SEM) was applied to verify the presence of bacteria and a fouling layer formation. For that a Zeiss Orion NanoFab (Carl Zeiss Microscopy, Peabody, MA, USA) was used. A beam of singly charged helium ions with an ion landing energy of 25kV and a beam current of about 1pA was scanned across the sample. Scanning electron imaging was done with an Everhart-Thornley type detector. Combined Fluorescence in Situ Hybridization (Combined-FISH), which integrates direct-gene FISH for ARG detection with catalyzed reporter deposition FISH for rRNA labelling of bacterial cells (Wang et al., 2026), was used to visualize potential plasmid deposition on the membrane surface. More details can be found in the SI7.

### 2.7. Calculations and data analysis

The *RE* (%) is expressed as the percentage of retention (Krzeminski et al., 2020):

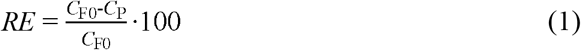

where *C*_F0_ (copies/µL) and *C*_P_ (copies/µL) are, respectively, the concentration of the respective ARGs in the feed at the start and their concentration in the permeate at the end of the filtration experiments. The overall mass balances were also calculated for all the filtration experiments to determine the dominant mechanisms for the removal (Slipko et al., 2019), including plasmid mass recovered in the retentate (Mr, rejection), plasmid mass recovered in the permeate (Mp, permeation), and the unrecovered plasmid mass attributed to membrane adsorption (Ma) (SI7). Additional calculations, including mass balance and fouling calculation in terms of membrane resistance can be found in SI8. The collected data were analyzed with the use of statistic tools available in RStudio with R (Posit Team 2024) (version 2024.09.0+375) (Allaire, 2012). All measurements were performed in triplicates and an arithmetic mean and standard deviation were calculated for every set of data. Differences in plasmid mass balance fractions (Mr, Ma, Mp) between the non-fouled and pre-fouled membrane were assessed using Wilcoxon rank-sum test (Wilcoxon, 1945).

## 3. Results and discussion

### 3.1. Validation of UF performance and biofouling development

To establish the experimental conditions for investigating plasmid retention under biofouling conditions, we first assessed the removal of ARB and the formation of a biofouling layer on the UF membrane.

Firstly, before investigating plasmid removal, the performance of the UF system was validated using *P. putida* KT2440 as a model bacterium. Synthetic effluent medium was spiked with *P. putida* KT2440, and bacterial concentrations in the feed and permeate were quantified by flow cytometry (Fig. 1(a)). ARB removal exceeded 99.99% during the first week and remained above 99.97% throughout the experiment, confirming that the 100 kDa MWCO UF membrane efficiently retained intact bacterial cells, in agreement with previous studies (Mota et al., 2022; Schwermer et al., 2018). These results established a robust baseline for the subsequent investigation of extracellular plasmid removal.

**Fig. 1.**
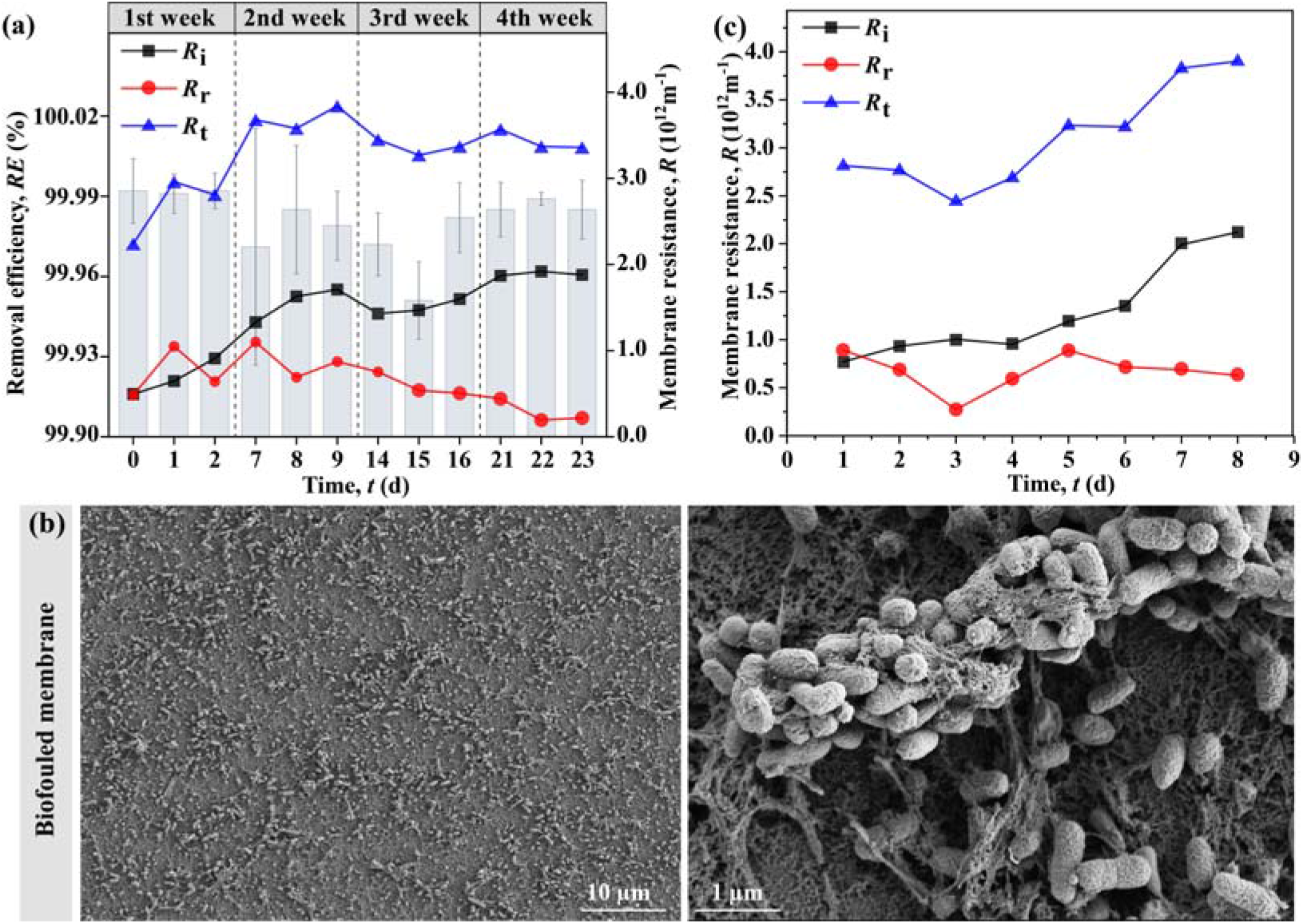
(a) Removal efficiency (*RE*, %) of ARB (*P. putida* KT2440) in the UF system (bars, mean±SD) and membrane resistances during the first pre-fouling experiment: reversible (*R*_r_) and irreversible (*R*_i_) membrane resistance, and total resistance (*R*_t_). (b) SEM analysis of fouled membrane. Scale bar is 10 µm and 1µm. (c) *R*_r_, *R*_i_ and *R*_t_ in the second pre-fouling experiment.

Next, we examined whether a biofouling layer developed on the membrane during 12 UF runs. The fouling was quantified by the total membrane resistance (*R*_t_), which is the sum of the intrinsic membrane resistance and the fouling resistance. The fouling resistance consists of a reversible membrane resistance (*R*_r_), which can be removed by hydraulic cleaning, and an irreversible membrane resistance (*R*_i_), which remains after hydraulic cleaning (SI8). During the consecutive filtration runs, an increase in *R*_t_ was observed (Fig. 1(a)). The increase was mainly driven by *R*_i_, whereas *R*_r_ did not increase, suggesting the gradual development of membrane fouling. To verify the nature of this fouling layer, the membrane was disassembled after completion of the pre-fouling experiment and analyzed by SEM (SI7). The membrane autopsy revealed a dense biofilm consisting of *P. putida* KT2440 cells embedded in extracellular polymeric substances (EPS) (Fig. 1(b)), confirming the successful establishment of a biofouled membrane. Such a biofilm forms an additional layer on the membrane surface that hinders water flow through the membrane, thereby increasing *R*_i_, consistent with previous reports describing biofilm formation during membrane filtration (Cheng and Hong, 2017; Zarei-Baygi et al., 2020).

### 3.2. Quantitative effects of biofouling on plasmid retention

#### 3.2.1. Biofouling alters plasmid mass partitioning during ultrafiltration

As the membrane used in the previous experiment had been removed from the cassette for SEM analysis, a new pre-fouled membrane was prepared for the subsequent plasmid retention experiments. To this end, an independent pre-fouling experiment was conducted with *P. putida KT2440* suspensions filtered over eight consecutive runs (Section 2.5). The progressive increase in *R*_i_ confirmed the development of membrane fouling (Fig. 1(c)). Seven consecutive filtration runs were performed using either an initially non-fouled membrane or the fouled membrane to determine how established biofouling affects the retention of the ARG-carrying plasmid R388. To elucidate the mechanisms governing plasmid removal, a mass balance analysis was performed for each filtration run. By quantifying the plasmid mass recovered in the medium and permeate and accounting for the unrecovered fraction associated with the membrane, the relative contributions of Mr, Mp, and Ma were determined (Fig. 2).

**Fig. 2.**
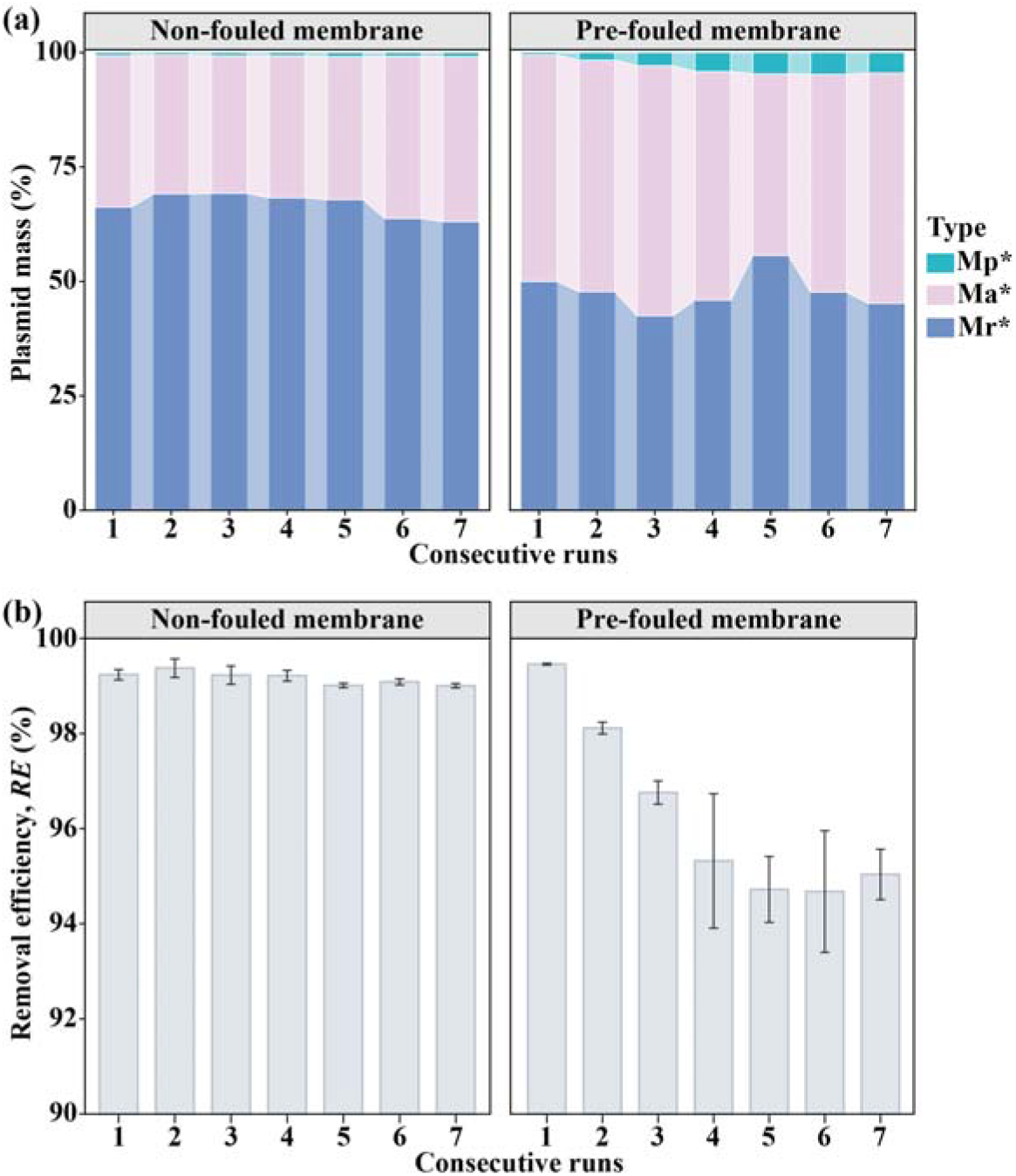
(a) Mass balance of plasmid R388 for the non-fouled and pre-fouled membrane across seven consecutive runs. The mass balance tracks the fate of the initial plasmid mass introduced into the system, partitioned into three fractions: plasmid mass recovered in the retentate (Mr, rejection), plasmid mass recovered in the permeate (Mp, permeation), and the unrecovered plasmid mass attributed to membrane adsorption (Ma). Asterisks indicate statistically significant differences between the non-fouled and pre-fouled membrane for each mass balance fraction (* paired Wilcoxon rank-sum test, Holm-adjusted *p* < 0.05). (b) Removal efficiency (*RE*, %) of R388 for the non-fouled and pre-fouled membrane across the same seven runs (mean±SD).

For the non-fouled membrane, the majority of the plasmid was consistently rejected (65−70%), while adsorption onto the membrane surface contributed approximately 30–35%, which indicated that membranes with a MWCO of 100 kDa could retain most of the plasmid introduced to the system, while only a minor fraction passed the membrane to reach the permeate (Fig. 2(a)). After the fifth filtration cycle, a slight increase in the plasmid concentration in the permeate was observed, and the *RE* gradually decreased to around 99.0% by the final run (Fig. 2(b)). However, this variation remained small and not statistically significant, indicating that the non-fouled membrane exhibited stable and reproducible performance throughout the consecutive filtrations.

In contrast, when a pre-fouled membrane was used the plasmid concentrations in the permeate increased significantly (Holm-adjusted *p* = 0.047) over the course of the experiment. While at the first filtration, *RE* was still at 99.4%, it already dropped below 99% in the second and below 95% in the last filtration (Fig. 2(b)). This increased plasmid number in the permeate is unlikely to result from a change in membrane pore size, as no higher transmembrane pressure was applied that could promote pore enlargement, and excessive membrane compaction was performed to prevent pore elongation. This indicates that the increase is related to the presence of the biofouling layer rather than to the membrane itself. Our results suggest that plasmids from previous filtrations may transfer to the permeate in subsequent runs, as shown by the gradual increase in permeate concentration.

Unlike the case of the non-fouled membrane, rejection was no longer the dominant removal mechanism. Instead, biofouling substantially altered the distribution of plasmids among the three mass balance fractions, with rejection decreasing to an average of 48% and membrane-associated adsorption increasing to 49% across the seven filtration runs, accompanied by a smaller increase in the permeate fraction. These findings suggest that biofouling changes plasmid transport through the membrane, shifting plasmid removal mechanisms from rejection towards membrane adsorption and permeation. The higher adsorption on the pre-fouled membrane is consistent with the plasmid–EPS interaction (Section 3.2.2), and a gradual increase in this interaction over the successive runs may allow more plasmids to pass through the pores.

#### 3.2.2. Microscopic evidence for biofilm-associated plasmid retention

After the seven consecutive R388 filtration runs, the initially non-fouled and pre-fouled membrane cassettes were disassembled, and the membrane surfaces were examined by HIM and Combined-FISH to determine how the two initial fouling conditions affected membrane-associated plasmid accumulation.

Observation of the unfouled membrane surface after seven runs using HIM revealed deposition of material originating from the synthetic effluent medium on top of the membrane, thereby covering the porous structure observed for a pristine membrane (Fig. 3(a) and (b)). Since there were no bacteria in the feed, no EPS-like structures were found, and also no change in the permeate flux was recorded throughout the consecutive runs. To verify that plasmids contributed to the deposited material, Combined-FISH was performed, targeting the *sul1* ARG located on the plasmid (Fig 3(d-e)). Similarly, with the HIM imaging results the Combined-FISH showed patchy distribution of the *sul1* gene and plasmids on the membrane surface (Fig. 3 (d)). These results are consistent with size exclusion contributing to plasmid retention (Section 3.2.1), in addition to membrane-associated adsorption. In addition, R388 was present predominantly in its native supercoiled conformation during these experiments. Its highly branched and sterically complex structure may have increased hydrodynamic resistance to pore passage, thereby favoring retention at the membrane under these non-fouled conditions (Rathore and Velayudhan, 2002).

**Fig. 3.**
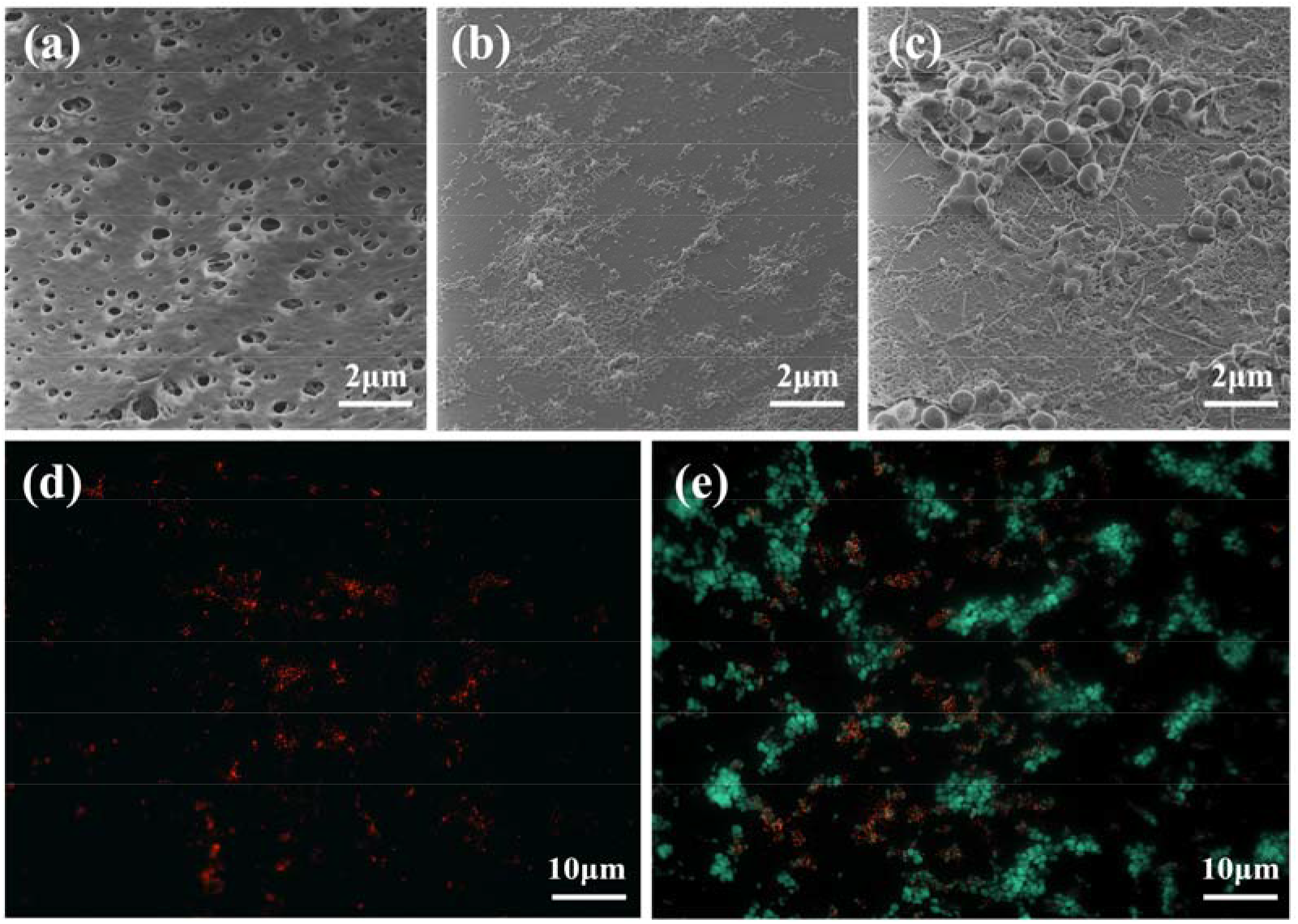
HIM micrographs of the (a) pristine, (b) non-fouled and (c) pre-fouled membranes with a scale bar of 2 µm, and (d) Combined-FISH for the non-fouled and (e) pre-fouled membranes. In figure d and e, the red dot-like signals are *sul1* gene signal, which is labeled by Alexa594 fluorochrome, whereas the green signals indicating *E. coli* cells that were labeled with Alexa488 fluorochrome.

In contrast, the pre-fouled membrane already contained a dense biofilm prior to the R388 filtration experiments (Section 3.1). After the seven filtration runs, the biofilm showed a high spatial heterogeneity with respect to EPS structures on the membrane surface (Fig. 3(c)). The areas with higher plasmid deposition coincided with regions of increased biofouling (Fig. 3(e)). This is most likely attributed to the high DNA adsorption capacity of EPS (Li et al., 2024) and electrostatic interactions between the negatively charged DNA and the biofilm matrix, providing many additional binding sites for plasmids compared to the pristine membrane surface. This biofilm-mediated adsorption is consistent with the pronounced increase in the adsorbed fraction observed in the mass balance for the pre-fouled membrane (Section 3.2.1). On this membrane, the adsorbed fraction (49%) became comparable to rejection (48%). In contrast, rejection remained the dominant mechanism on the non-fouled membrane. Plasmids may therefore have accumulated in the fouling layer, which could have increased the plasmid concentration near the pores in later runs. Plasmids that remain at the membrane−biofilm interface close to the pores have a longer residence time and are more likely to pass through them, as the converging flow fields near the pores may promote their deformation (Arkhangelsky et al., 2011; Latulippe and Zydney, 2011). In membrane-based DNA purification, plasmids have been described to stretch into a “snake-like” shape to enter membrane pores and to return to their normal form after passing through (Arkhangelsky et al., 2011). Additionally, although fouling may alter local transport conditions, there is no direct evidence that such effects contributed to the observed changes in plasmid permeation. Our results highlight the limitations of assessing plasmid *RE* based on single filtration runs with clean membranes. While such experiments provide information on initial membrane performance, they fail to capture the effects of biofilm formation and development on plasmid retention during wastewater treatment. In our study, the presence of a biofouling layer altered plasmid retention, with *RE* declining over consecutive filtration runs on pre-fouled membranes. These findings demonstrate that biofouling is not merely an operational challenge but an important factor influencing the retention and permeation of extracellular plasmids. Consequently, *RE* determined using clean membranes alone may not adequately represent membrane performance under wastewater treatment conditions, emphasizing the need to account for biofilm development when evaluating membrane-based removal of extracellular ARGs.

### 3.3. Intrinsic plasmid properties govern retention and permeation

Having established that membrane biofouling alters the retention of the model plasmid R388, we next performed a separate set of filtration experiments to determine whether intrinsic plasmid properties also affect UF behavior. To isolate the effects of plasmid size and conformation from those of established biofouling, plasmids of different sizes were tested in linear and supercoiled forms using non-fouled membranes.

In all cases, no clear size dependence was observed, as RSF1010 (8.7 kbp) exhibited the lowest *RE* values (86.4±4%and 98.5±0.3% for linear and supercoiled, respectively), regardless of its conformation (Fig. 4(a)). Actually, among the studies on DNA purification, there is no agreement regarding the effect of plasmid size on the *RE* of UF membranes. For example, Latulippe and Zydney (2011) tested UF for plasmids ranging in size from 3 to 17 kbp and found that permeation was independent of plasmid molecular size. Opposite, Li et al. (2016) found that *RE* increased with plasmid size (3, 9.8 and 16.8 kbp), although those differences could only be detected above certain flux values. For a wastewater context, no studies have been found for the comparison of different plasmid sizes. In this work, *RE* did not follow a monotonic trend with plasmid size within the 3.3–12.3 kbp range, suggesting that plasmid-specific characteristics other than size may contribute to the removal differences. Unlike plasmid size, plasmid conformation had a clear effect on removal. Supercoiled plasmids showed consistently higher removal rates (mean values of 98.8−99.3%) compared to linearized plasmids (mean values of 89.3−95.0%). This indicates that conformation had a more pronounced effect on *RE* than plasmid size within the tested range, particularly for linear plasmids.

**Fig. 4.**
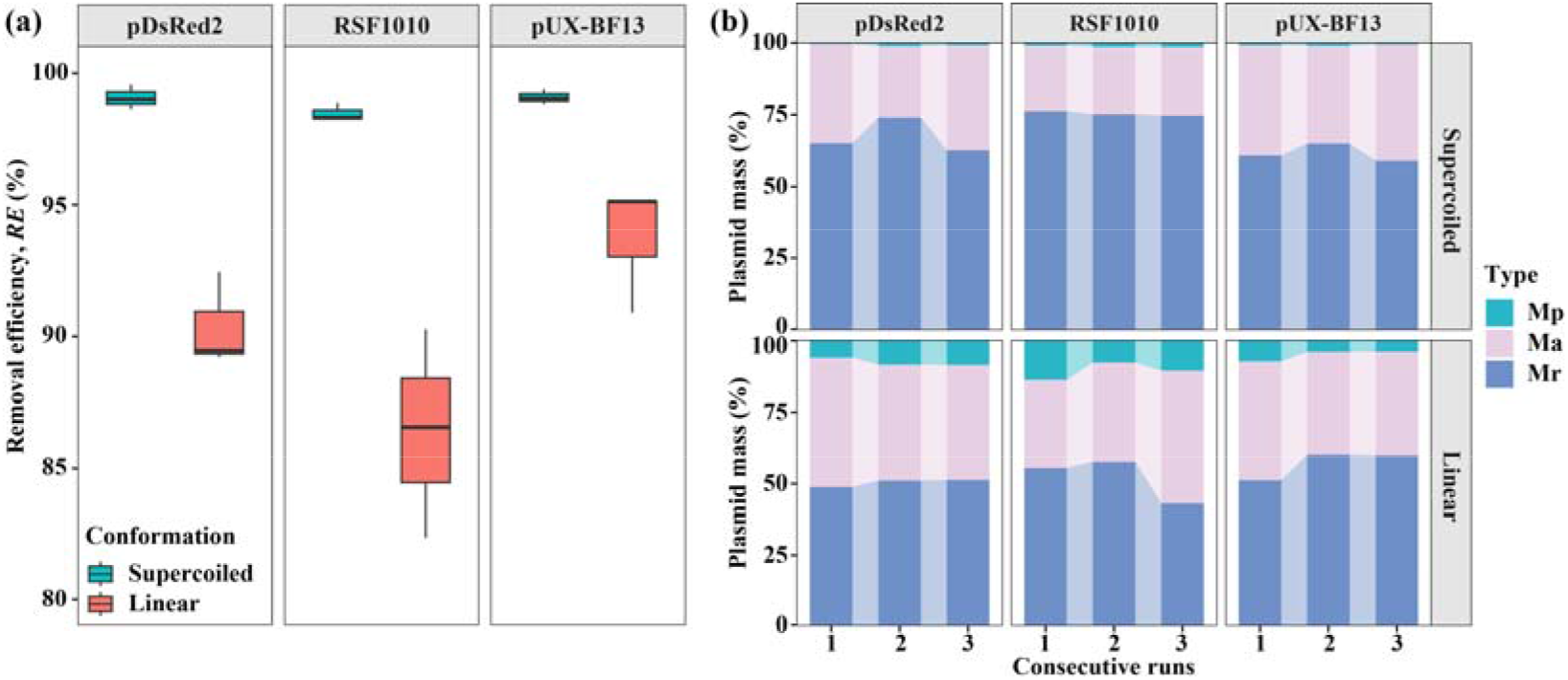
Effects of plasmid size and conformation on UF retention. (a) Removal efficiency (*RE*, %) of the linear and supercoiled isoforms for different sized plasmids. (b) Mass balance of different plasmid concentrations in which Mp, Ma and Mr is plasmid masses from the permeate, absorption, and total rejection.

In terms of mass distribution, all supercoiled plasmids exhibited consistently high rejection (Mr, 58.9–76.2%) and a substantial adsorption (Ma, 22.9–40.6%), whereas the permeate fraction was negligible (Mp, 0.3–1.4%) (Fig. 4(b)). In contrast, linear plasmids showed a reduced Mr (43.1–60.1%) and a markedly increased Mp (3.9–13.8%), approximately 4–9 times that of their supercoiled counterparts, which accounts for their lower *RE* values. Moreover, a small difference was observed in rejection for the large linear plasmids (i.e., RSF1010 and pUX-BF13) compared with the smaller one (i.e., pDsRed2), indicating that there is no clear relationship between rejection and plasmid size, which is consistent with the absence of a clear size dependence of *RE*. Among linearized plasmids, RSF1010 exhibited the lowest *RE* (Fig. 4(a)). The lower permeation of supercoiled plasmids may be related to the branched structure resulting from the under-twisting of the DNA whereas linear plasmids, not having such branched structures, would be more likely to pass through the membrane pores (Li et al., 2016). Moreover, linear plasmids lack the torsional constraint of supercoiled plasmids and are expected to adopt less complex, more extended conformations. They may therefore be more easily elongated by the flow field, enabling “snake-like” movement through local pore openings (Latulippe and Zydney, 2009), which may contribute to their higher permeation. However, the transfer of large plasmids through relatively small pore-size membranes is strongly limited, as these long and highly entangled molecules are more easily trapped or “jammed’’ at pore entrances (Arkhangelsky et al., 2011). This may help explain why the largest plasmid pUX-BF13 showed the lowest permeate fraction among the linear plasmids (5.0±1.9% vs. 7.6±1.5% for pDsRed2 and 10.7±3.0% for RSF1010). Moreover, DNA molecules can form knots limiting their ability to elongate under flow, which may further hinder their passage through membrane pores. Differences in the tendency of plasmids to form such knots may therefore have contributed to variations in their membrane retention and permeation observed in this study (Rosa et al., 2012). Therefore, the permeation of plasmids through the membrane appears to be shaped by several interacting factors, including plasmid-specific characteristics.

In this work, the effect of intrinsic plasmid properties was investigated separately using initially non-fouled membranes. However, for wastewater applications, UF membranes inevitably operate under fouling-prone conditions, and thus plasmids may interact with the developing fouling layer that facilitates either adsorption or partial transmission. As a result, the overall retention of plasmid DNA in wastewater treatment may not be predicted only from intrinsic molecular size or conformation and may also depend on membrane fouling, the water matrix and surface interactions. For instance, the presence of colloids in the water has been reported to promote the retention of ARGs (Breazeal et al., 2013), whereas higher ionic strength has been reported to results in greater compaction of DNA molecules (Latulippe and Zydney, 2008) for which further research should assess plasmids permeation through fouled membranes under different effluent-like water qualities. Based on the experimental findings, Fig. 5 presents a conceptual overview of how membrane fouling and plasmid-intrinsic properties, including size and conformation (linear versus supercoiled), influence plasmid retention during UF.

**Fig. 5.**
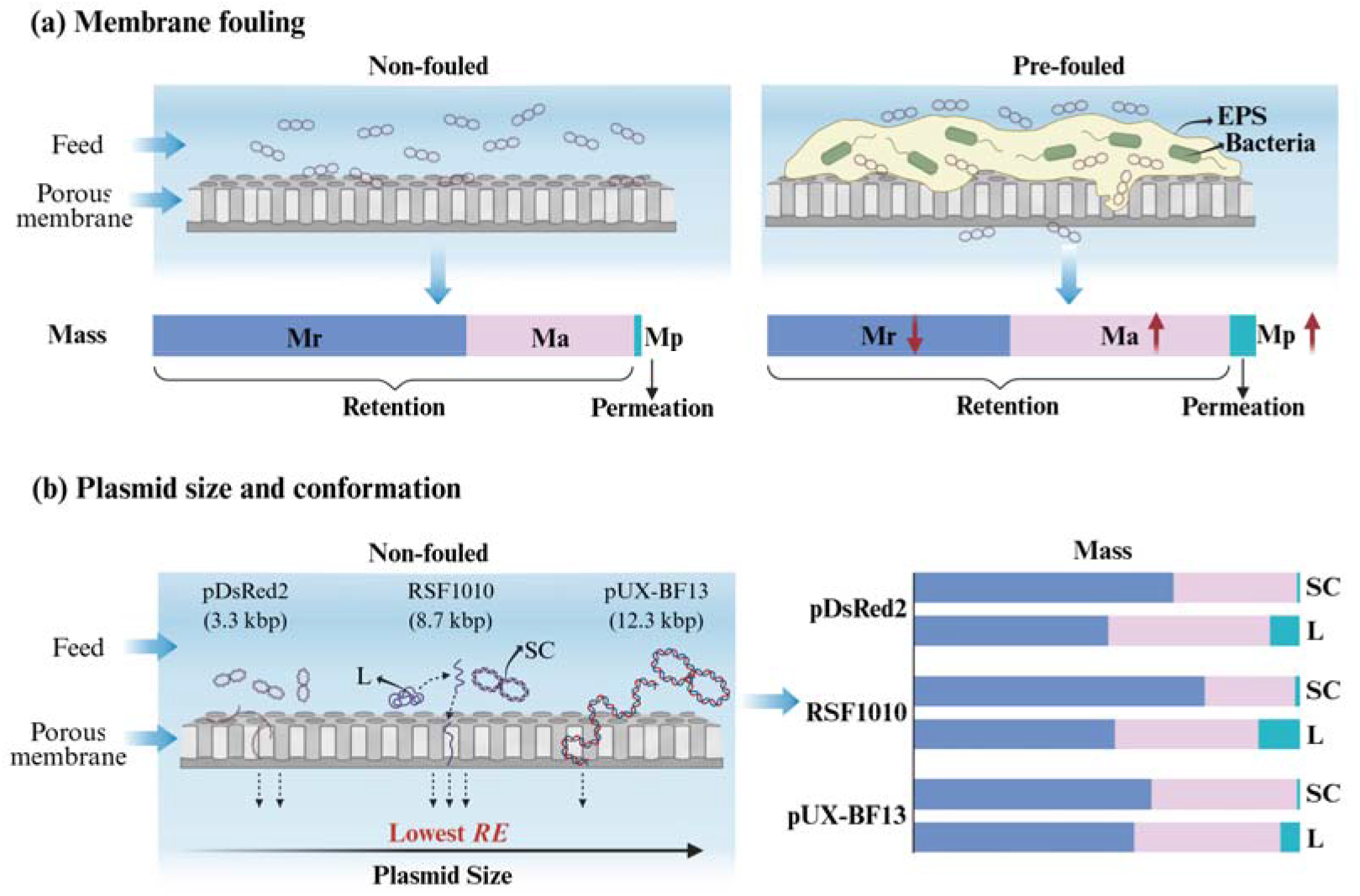
Conceptual summary of the fate of plasmids during UF. (a) Effect of membrane fouling: Distribution of the introduced R388 plasmid mass among rejection (Mr), adsorption (Ma) and permeation (Mp) for non-fouled and pre-fouled membranes. (b) Effect of plasmid size and conformation on non-fouled membranes: mass partitioning of supercoiled (SC) and linear (L) plasmids of 3.3, 8.7 and 12.3 kbp. Bars show mean values (*n* = 3 runs for (b); 7 runs for (a)). Created with BioRender (agreement number RQFU6E5JVN).

## 4. Environmental implications

With the revised Urban Wastewater Treatment Directive extending requirements for advanced treatment and the monitoring of antimicrobial resistance in wastewater, additional treatment for the release of ARGs are increasingly relevant. Our findings highlight the potential of UF as a tertiary treatment barrier against extracellular plasmid-associated ARGs from WWTPs. A 100 kDa MWCO membrane achieved *RE* of approximately 99% for supercoiled plasmids and 82−92% for linear plasmids, suggesting that low-pressure UF may improve ARG retention without more restrictive processes such as nanofiltration.

However, UF effectiveness depends on more than nominal membrane pore size. Membrane biofouling shifted plasmids from rejection to adsorption and permeation, lowering overall *RE*. This indicates that biofilm accumulation should not automatically be considered an additional protective barrier against extracellular ARG release. Future research should therefore investigate how fouling development, cleaning strategies, and operational conditions affect plasmid retention over extended periods and whether sequential membrane configurations could provide more stable removal.

Importantly, our findings also have implications for how membrane-based ARG removal is evaluated. Supercoiled plasmids were consistently retained more efficiently than their linear counterparts, while plasmid size alone was not a reliable predictor of *RE*. Because extracellular DNA in treated wastewater may occur in different conformations and fragment sizes, experiments relying exclusively on intact supercoiled plasmids may overestimate the retention of the broader extracellular DNA pool. Future assessments should therefore consider a wider range of DNA structures and include mass balance approaches that distinguish rejection, adsorption, and permeation rather than relying solely on overall *RE*. The linearized plasmids used here were generated artificially from circular plasmids and differ from naturally occurring linear plasmids, which can replicate autonomously, so whether the higher permeation of linearized DNA translates into a higher potential for HGT remains to be shown.

Additionally, from the practical and economic perspective, UF is a simple, highly automated process with a low footprint (Chew et al., 2016). Its energy demand is higher than that of conventional tertiary treatments such as ultraviolet disinfection or ozonation, but it achieves higher effluent quality. As a low-pressure process, UF has lower energy and operating costs than nanofiltration or reverse osmosis (Jiménez-Benítez et al., 2024). Costs depend strongly on membrane fouling, which increases energy demand and cleaning requirements and shortens the membrane lifespan (Al Aani et al., 2020). Because biofouling reduced plasmid retention and irreversible resistance increased despite hydraulic cleaning, periodic cleaning or membrane replacement will probably be needed to maintain the *RE*. UF has been demonstrated at demonstration and semi-industrial scale for municipal wastewater (Jiménez-Benítez et al., 2024), but pilot-scale tests with real effluents are needed to confirm scalability.

Together, effective control of extracellular ARG discharge requires consideration of both membrane operating conditions and the structural characteristics of the DNA being retained. Incorporating these factors into the design and evaluation of tertiary treatment technologies could help establish more realistic performance criteria and support the development of strategies to limit the environmental dissemination of ARG.

## Acknowledgement

W.X.C. thanks the China Scholarship Council (CSC202306090243) for a 2-year grant to study abroad. F.R.S. acknowledges funding from the European Union’s Horizon 2020 research and innovation programme under the Marie Skłodowska–Curie grant agreement No. 101022115. N.M. acknowledges funding by the Novo Nordisk Foundation through Grant NNF22OC0071609 ReFuel. A.W. is grateful for funding from the *Deutsche Forschungsgemeinschaft* (DFG, German Research Foundation), project number *530107960*. The authors are grateful for using the helium-ion microscope at ProVIS–Centre for Chemical Microscopy at the Helmholtz–Centre for Environmental Research, Leipzig, which is supported by European Regional Development Funds (EFRE–Europe funds Saxony), and the Helmholtz Association. The content of this article reflects only the authors’ views and the Research Executive Agency is not responsible for any use that may be made of the information it contains.

## Author contributions

**Wenxuan Chen**: Writing − original draft, Formal analysis, Visualization. **Fátima Rojas Serrano**: Writing – review & editing, Methodology, Data curation, Investigation. **René Kallies**: Writing – review & editing, Investigation. **Susann Müller**: Writing – review & editing, Investigation. **Niculina Musat**: Writing – review & editing, Investigation. **Florian Schattenberg**: Writing – review & editing, Investigation. **Matthias Schmidt**: Writing – review & editing, Investigation. Hendrik Seifert: Writing – review & editing, Investigation. **Gangan Wang**: Writing – review & editing, Investigation. **Anja Worrich**: Writing – review & editing, Supervision, Conceptualization.

