## Supporting information for "Membrane Biofouling and Plasmid Conformation Shape the Retention of Antibiotic Resistance Gene-Carrying Plasmids during Ultrafiltration"

**List of contents:**

SI1. Synthetic effluent and feed water preparation

SI2. Plasmids extraction and linearization

SI3. Experimental facility

SI4. Quantitative real time PCR

SI5. Fouling pre-experiment with bacteria

SI6. Flow cytometric measurement of cell numbers

SI7. Fouling visualization with HIM/SEM and Combined-FISH

SI8. Additional calculations

*Reference style in supporting files is different than Journal format.

### SI1. Synthetic effluent and feed water preparation

The trace elements and vitamins were added in a ratio of 12.5 mL/L to the synthetic effluent composition, and the composition of the respective solutions can be found in Table S1 and Table S2.

**Table S1.** Composition of trace elements solution.

| **Component** | **Concentration (g/L)** |
|---|---|
| Nitrilotriacetic acid | 1.50 |
| $MgSO_4 \cdot 7H_2O$ | 3.00 |
| $MnSO_4 \cdot H_2O$ | 0.50 |
| NaCl | 1.00 |
| $FeSO_4 \cdot 7H_2O$ | 0.10 |
| $CoSO_4 \cdot 7H_2O$ | 0.18 |
| $CaCl_2 \cdot 2H_2O$ | 0.10 |
| $ZnSO_4 \cdot 7H_2O$ | 0.18 |
| $CuSO_4 \cdot 5H_2O$ | 0.01 |
| $KAI(SO_4)_2 \cdot 12H_2O$ | 0.02 |
| $H_3BO_3$ | 0.01 |
| $Na_2MoO_4 \cdot 2H_2O$ | 0.01 |
| $NiCl_2 \cdot 6H_2O$ | 0.03 |
| $Na_2SeO_3 \cdot 5H2O$ | 0.30 |
| $Na_2WO_4 \cdot 2H_2O$ | 0.40 |

**Table S2.** Composition of vitamin solutions.

| **Component** | **Concentration (g/L)** |
|---|---|
| Biotin (H) | 2.00 |
| Folic acid ($B_2$) | 2.00 |
| Pyridoxin-HCl ($B_6$) | 10.00 |
| Thiamin-HCl ($B_1$) | 5.00 |
| Riboflavin ($B_2$) | 5.00 |
| Nicotinic acid ($B_3$) | 5.00 |
| Calcium pantothenate | 5.00 |
| Cyanocobalamin $B_{12}$ | 0.10 |
| p-Aminobenzoic acid | 5.00 |
| Lipoic acid | 5.00 |

The current European Urban Wastewater Treatment Directive (Council Directive 91/271/EEC)[1] establishes the COD of 125 mg/L $O_2$ as maximum limit for wastewater discharge of secondary effluents. To guarantee the feed water used within the experiments fulfilled the European requirements for secondary effluents, the COD was measured before and after the addition of the plasmids with NANOCOLOR® CSB 160" test (Macherey-Nagel, GmbH & Co. KG, Germany), obtaining 61 mg $O_2$/L for the synthetic effluent without the plasmids and up to 80 mg $O_2$/L values after the addition of the plasmids, for which we can ensure the feed water for the experiments was comparable to a real secondary effluent.

### SI2. Plasmids extraction and linearization

Two different plasmid extraction/purification kits were used because the amount of plasmid DNA required differed for these experiments. Nucleospin Plasmid® (Macherey-Nagel GmbH & Co. KG, Germany) was used for the extraction of R388 plasmid, used in Part I, whereas NucleoBond® PC 100 Midi Kit (Macherey-Nagel GmbH & Co. KG, Germany) was used for the extraction of pDsRed2, RSF1010 and pUX-BF13 plasmids, tested in Part II. All the plasmids are low-copy and thus, the corresponding procedures indicated by the manufacturer were followed, but the extraction yields for the different plasmids were quite variable. After the extractions, the respective plasmids solutions were mixed in low-binding Eppendorf tubes (1.5 mL) and the concentrations of the resulting mother solutions were measured with fluorescence by "Invitrogen™ Qubit™ dsDNA BR-Assay-Kit" (Thermo Fisher Scientific Inc., USA).

All plasmid sequences were checked in silico for respective single cut restriction sites with NEB cutter. Although only the supercoiled form of R388 plasmid was used within the experiments in Part I, a little volume of the extracted plasmid was linearized with EcoRI enzyme (12 units/µL, Promega Corporation) to compare the linear plasmid to the one after extraction, expected to be supercoiled. Electrophoresis with 0.8% agarose gel[2-3] was performed (gel loading dye and Quick-Load® 1 kb Extend DNA ladder by New England Biolabs Inc., 100 V, 75 min, 15 min incubation with ethidium bromide). As shown in Fig. S1, only one band could be detected for the digested plasmid (Lane 8) between 20 and 48.5 kb, matching the plasmid size (33,926 bp). The lanes filled with volumes of undigested plasmid solution (Lanes 10 to 12) also matched the plasmid size but the bands were blurry and slightly lower than that of the digested plasmid, as usual for supercoiled plasmids. There were additional bands for the undigested plasmid lanes far beyond the size of the ladder, probably corresponding to genomic DNA flushed during plasmids extraction.

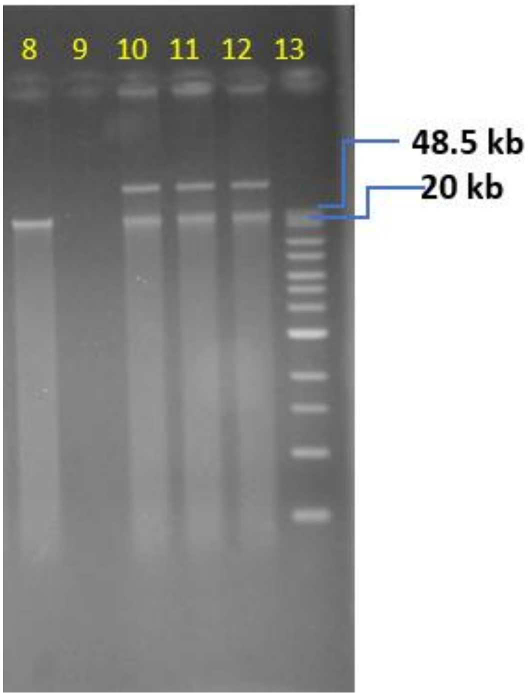


**Fig. S1**. Plasmid R388 with and without digestion. Lane 8 = Digested plasmid with EcoRI, Lane 9 = Empty; Lanes 10-12 = Undigested plasmid; Lane 13 = Ladder.

The plasmids used within Part II were linearized by digestion with respective restriction enzymes. EcoRI (12 units/µL, Promega Corporation, USA) was used for pDsRed2 and RSF1010 whilst SphI-HF (20 units/µL, New England Biolabs Inc., USA) was used for pUX-BF13. The choice of enzymes was based on the restriction sites of the respective plasmids, with the criterion of cutting only one site and without affecting the resistance gene of interest[4-6]. To linearize plasmids pDsRed2 and RSF1010, deionized water, Restriction Enzyme 10X Buffer, acetylated bovine serum albumin (BSA) (10 µg/µl) and EcoRI (12 units/µL) were used according to the manufacturer´s protocol (Promega Corporation, USA). To linearize plasmid pUX-BF13, deionized water, rCutSmart buffer, pUX-BF13, and SphI-HF (20 units/µL) were added as described in the supplier´s protocol (New England Biolabs Inc., USA).

The complete digestion of the plasmids was also verified with gel electrophoresis (same procedure as with R388 plasmid), as shown below.

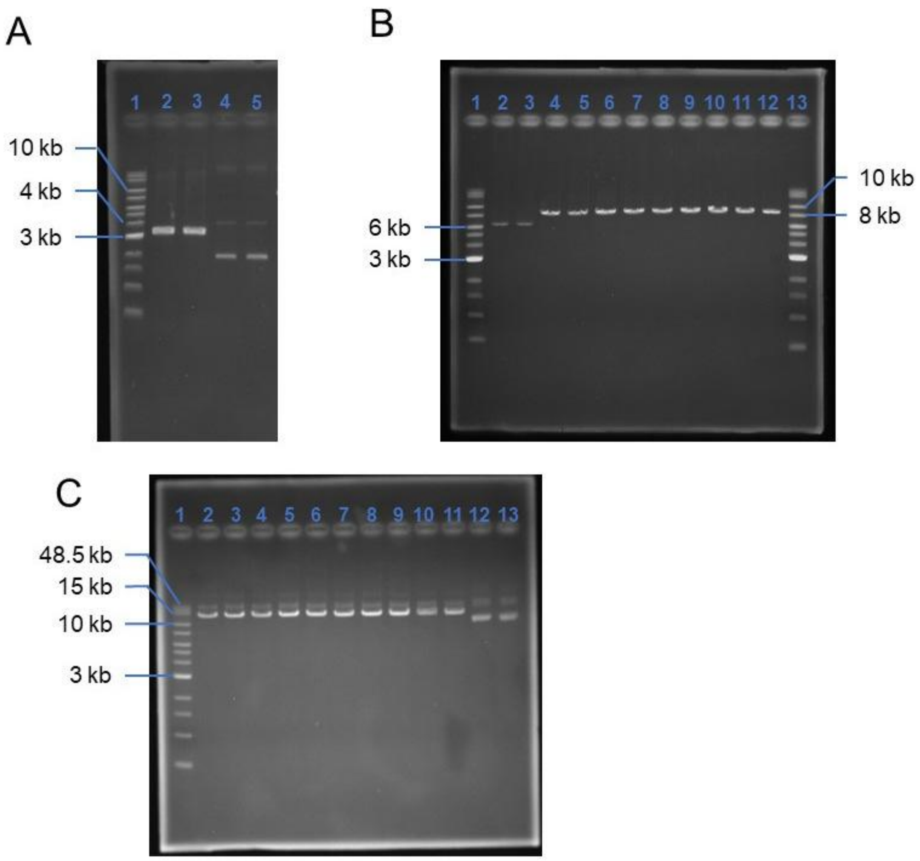


**Fig. S2**: Linearization of plasmids pDsRed2, RSF1010, and pUX-BF13. **A**, Linearization of pDsRed2 with restriction enzyme EcoRI. Lane 1 = DNA ladder, lane 2 and 3 = linearized pDsRed2, lane 4 and 5 = non-linearized pDsRed2. **B**, Linearization of RSF1010 with restriction enzyme EcoRI. Lane 1 and 13 = DNA ladder, lane 2 and 3 = non-linearized RSF1010, lane 4 – 12 = linearized RSF1010. **C**, Linearization of pUX-BF13 with restriction enzyme SphI-HF. Lane 1 = DNA ladder, lane 2 – 11 = linearized pUX-BF13, lane 12 and 13 = non-linearized pUX-BF13.

Fig. S2A shows that linearized pDsRed2 (3,311 bp) migrated as a strong band slightly above 3 kb, with no additional bands detectable, whereas non-linearized pDsRed2 showed a strong band below 3 kb and a weaker band at approximately 4 kb.

In Fig. S2B, non-linearized RSF1010 (8,684 bp) showed a single band near 6 kb, whereas linearized RSF1010 migrated as a single band slightly above 8 kb, consistent with its expected size. Similarly, linearized pUX-BF13 (12,256 bp) showed a strong band between 10 and 15 kb, whereas non-linearized pUX-BF13 migrated predominantly near 10 kb (Fig. S2C). Additional bands in non-linearized plasmid preparations have also been reported in previous studies[7,8] and may represent different plasmid conformations, such as nicked or open-circular DNA resulting from single-strand breaks. Overall, the observed differences in electrophoretic mobility between linearized and non-linearized plasmids support successful linearization, although the presence of minor fractions of other plasmid conformations cannot be excluded.

The concentrations of the linearized plasmids were measured with Invitrogen™Qubit™ dsDNA BR-Assay-Kit (Thermo Fisher Scientific Inc., USA) and as the reference to recalculate respective supercoiled plasmids concentrations in accordance with the digestion reactions. Removal efficiencies were calculated from plasmid copy numbers measured by qPCR in the feed and permeate and are therefore independent of these initial concentration estimates. All stock plasmids solutions were stored at 4°C until further use.

### SI3. Experimental facility

The set-up used for the ultrafiltration experiments as shown in Fig. S3.

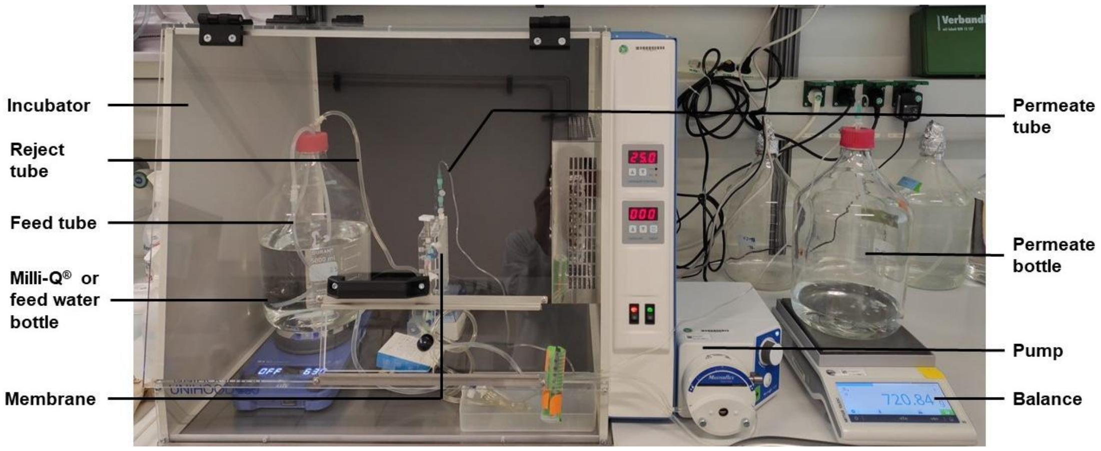


**Fig. S3**. Experimental facility used for the ultrafiltration experiments.

The membrane cassette (Vivaflow® 200 Laboratory Cross Flow Cassette, VF20P4, Sartorius AG, Germany) together with the feed water inlet and reject and permeate outlets are shown in Fig. S4. The membrane material was polyether sulfone and had a pore size of 100 kDa. Feed water inlet and reject outlet are on the right side whereas permeate outlet is on top.

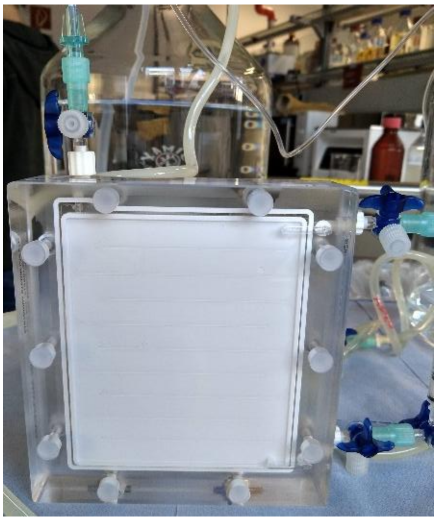

**Fig. S4**. Vivaflow 200 Laboratory Cross Flow Cassette (VF20P4, Sartorius AG, Germany).

### SI4. Quantitative real time PCR

The qPCR protocol comprised a denaturation step at 98 ºC for 2 min, followed by 40 cycles of amplification, as detailed in Table S3. The primers used are included in Table S4.

**Table S3**. Amplification step for the respective target genes.

| Target gene | Amplification step | Temperature, ºC | Time, s |
|---|---|---|---|
| *sul1 /sul2* | Denaturation | 98 | 5 |
| | Annealing/Extension + Plate read | 55 | 10 |
| *blaTEM* | Denaturation | 98 | 10 |
| | Annealing/Extension + Plate read | 60 | 30 |

**Table S4**. Primers used for qPCR.

| Target gene | Forward primer 5'-3' | Reverse primer 5'-3' |
|---|---|---|
| *sul1* | TGTCGAACCTTCAAAAGCTG | TGGACCCAGATCCTTTACAG |
| *blaTEM* | ATGTAACTCGCCTTGATCGT | CAATGATACCGCGAGACCC |
| *sul2* | CGGCTGCGCTTCGATT | CGCGCGCAGAAAGGATT |

Before the analyses of the feed water and permeate samples resulting from the ultrafiltration experiments, five independent standard curves were obtained for each target gene. Each standard curve was obtained from eight concentrations of target gene, ranging from 4 to $4 \times 10^7$ copies per µL of template, each measured in triplicate, which is above the minimum number of replicates suggested by Kralik et al. (2017)[9]. All five curves were included, and an average standard curve was calculated from the mean quantification cycle ($C$q)[9] values of the five curves for each concentration, so that all samples could be quantified with the same standard curve. Only average standard curves with an efficiency between 90% and 110% and $R^2 \geq 0.99$ were accepted. The amplicons used and the resulting average standard curves can be found in Table S5 and S6, respectively.

**Table S5**. Amplicons used for the standard curves.

| Target gene | Product size (bp) | Concentration (ng/µL) | Concentration (copies/µL) |
|---|---|---|---|
| *sul1* | 112 | 10.99 | 9.09E+10 |
| *blaTEM* | 267 | 44.3 | 1.54E+11 |
| *sul2* | 66 | 7.81 | 1.10E+11 |

**Table S6**. Average standard curves.

| Target gene | Slope | Intercept | $R^2$ | Efficiency (%) |
|---|---|---|---|---|
| *sul1* | -3.565 | 38.336 | 0.999 | 90.77 |
| *blaTEM* | -3.549 | 37.211 | 0.999 | 91.32 |
| *sul2* | -3.569 | 39.828 | 0.999 | 90.64 |

The limit of detection (LoD) and limit of quantification (LoQ) were calculated for each target gene with at least 10 replicates per standard concentration. The LoD was defined as the lowest concentration for which more than 95% positive signals were obtained. The LoQ was evaluated as the lowest standard concentration giving a coefficient of variation of less than 25%, the latter being defined as the ratio of the standard deviation of the total copy number to the mean of the total copy number for each standard concentration. The corresponding LoD and LoQ values are shown in Table S7.

**Table S7**. LoD and LoQ for the target genes used.

| Target gene | LoD (copies/µL) | LoQ (copies/µL) |
|---|---|---|
| *sul1* | 17.04 | 38.97 |
| *blaTEM* | 38.77 | 342.00 |
| *sul2* | 1.58 | 5.17 |

The analyses of feed water revealed that the plasmid concentrations measured by qPCR (expressed in copies/µL) did not match with the expected values calculated by Eq.S1 and were significantly lower than expected for all supercoiled plasmids (Table

S8). In contrast, qPCR concentrations for linear plasmids were in the expected order of magnitude, suggesting that plasmid conformation affects DNA quantification by qPCR in a similar way to that observed for the fluorometric methods previously described (SI2). However, the ratio between the concentrations measured and the expected ones was always constant for every plasmid (Table S8), for which the removal efficiencies, the main focus of the work, would not be affected.

$$N\ (\text{copies}/\mu\text{L}) = C \times N_A/(L \times 660 \times 10^9) \times 10^{-6} \quad \text{(S1)}$$

where $C$ is the DNA mass concentration in the feed water (ng/L), $N_A$ is Avogadro's constant ($6.022\times10^{23}$ $\text{mol}^{-1}$), $L$ is the plasmid size (bp), 650 g/mol is the average molecular weight of one base pair, $10^9$ converts ng to g (the denominator is expressed in ng/mol), and $10^{-6}$ converts L to μL.

Several papers in the literature have pointed out the differences in qPCR calibration curves obtained with equal concentrations of the same plasmid in circular and linear conformations. According to Hou et al. (2010)[10], equimolar standard concentrations of supercoiled and linear plasmids, resulted in significantly higher *C*q for the linear plasmid standards. Because our qPCR standards were linear PCR amplicons (Table S5), a conformation-dependent amplification bias would lead to an underestimation of supercoiled plasmid concentrations. This could explain why supercoiled plasmids showed lower qPCR concentrations than expected. In contrast, linearized plasmids and linear PCR amplicons provided similar quantifications. Other authors such as Martinez-Fernandez et al. (2021)[11] reached similar conclusions and hypothesized that linearized DNA may be easier for the primers to bind compared to supercoiled plasmids. Conversely, Werling et al. (2015)[12] reported no significant effect of the plasmid conformation on DNA quantification.

**Table S8.** Expected and measured (qPCR) plasmid concentrations in the feed water

| Plasmid (conformation) | Size (bp) | Concentration in feed (ng/L) | Expected concentrations (copies/µL) | Measured concentrations (copies/µL) | | Measured/expected ratio (mean ± SD) |
|---|---|---|---|---|---|---|
| | | | | Mean | SD | |
| pDsRed2 (supercoiled) | 3311 | 10993 | 3.08E+06 | 2.91E+05 | 2.55E+04 | 0.10 ±0.01 |
| pDsRed2 (linear) | 3311 | 12589 | 3.52E+06 | 2.58E+06 | 2.26E+05 | 0.75 ±0.07 |
| RSF1010 (supercoiled) | 8684 | 5233 | 5.58E+05 | 8.53E+04 | 3.80E+03 | 0.16 ±0.01 |
| RSF1010 (linear) | 8684 | 6279 | 6.70E+05 | 4.16E+05 | 2.37E+04 | 0.64 ±0.04 |
| pUX-BF13 (supercoiled) | 12256 | 9711 | 7.34E+05 | 3.45E+05 | 1.23E+04 | 0.49 ±0.02 |
| pUX-BF13 (linear) | 12256 | 13950 | 1.06E+06 | 9.51E+05 | 3.51E+04 | 0.93 ±0.03 |

**SI5. Fouling pre-experiment with bacteria**

The fouling pre-experiment consisted in using a new membrane to filter a bacterial solution containing the model bacterium *P. putida* K2440 in a synthetic effluent medium over 4 weeks, with 3 UF runs per week (Fig. S5). This way, the membrane surface would be covered with a biofouling layer before the second batch of plasmid experiments in Part I was performed (section 2.5).

Precultures with *P. putida* KT2440 were prepared one day before each run by inoculating a small scoop from the agar-plate into a 120 mL R2A medium. The flask with the inoculum was placed in a shaking incubator (SM-30, Edmund Bühler GmbH, Germany) at room temperature (RT) overnight. The day after, R2A medium was replaced by synthetic effluent medium for which the preculture was centrifuged (6000 g, RT, 10 min, HERMLE Z326 K, Hermle) with the supernatant being discarded. The cell pellets were then resuspended in synthetic effluent medium using a vortex mixer (Vortex-Genie™ 2, Scientific Industries, Inc. New York, USA). After that, the concentration of the cell suspension was quantified with "QUANTOM™ Total Cell Staining Kit" (Logos biosystems, South Korea) according to the supplier's protocol (https://www.labtech.com/product/quantom-tx). This method was used for a rapid estimate of the feed concentration (counting range of the kit: $2\times10^5 - 1\times10^9$ cells/mL), whereas the lower concentrations in the permeate were quantified by flow cytometry (SI6). Once the concentration of the preculture was established, a certain volume was added to autoclaved synthetic effluent medium so that the final cell concentration was $2.0\times10^7$ cells/mL.

Before the UF experiments were performed, a growth curve in synthetic effluent and RT was performed in triplicates to confirm there would be no bacterial growth during the UF experiments for the model strain. The synthetic effluent has a low organic content (COD of 61 mg $O_2$/L; SI1), and the lag phase duration turned out to be 4 h, longer than the UF experiments, proving that the bacteria in the feed would either be retained by the membrane or reach the permeate but would not replicate in the course of the experiments.

The UF tests were performed with the same experimental set-up and conditions described in Material and Methods section 2.3. The membrane cassette remained inside the incubator at RT between consecutive tests so as to favor the formation of a biofilm on the UF membrane surface. Before every new test was performed, fresh feed was prepared as described above and permeate bottles and tubing were replaced by sterile ones. Additionally, blank filtrations with 5 L autoclaved Milli-Q® water were performed before and after each ultrafiltration batch. Sterile water was filtered through the system between consecutive experimental runs for two purposes. First, it flushed residual feed and non-attached bacteria from the membrane cassette and tubing, ensuring that each run started with fresh feed under comparable conditions. Second, this gentle rinsing step was intended to limit excessive fouling and prevent rapid membrane clogging while preserving the existing biofilm, allowing it to develop further over successive filtration runs. Once the third weekly run had been performed, the experimental facility was disassembled inside the sterile bench, the membrane inlet and outlets were closed with sterile plugs and placed in the fridge, remaining at 4 ºC until the week after, when the new tests were conducted. This procedure allowed controlling the progressive development of the biofilm over the 4 weeks period. After the pre-fouling experiment, the facility was disassembled and the membrane remained at 4ºC until the week after.

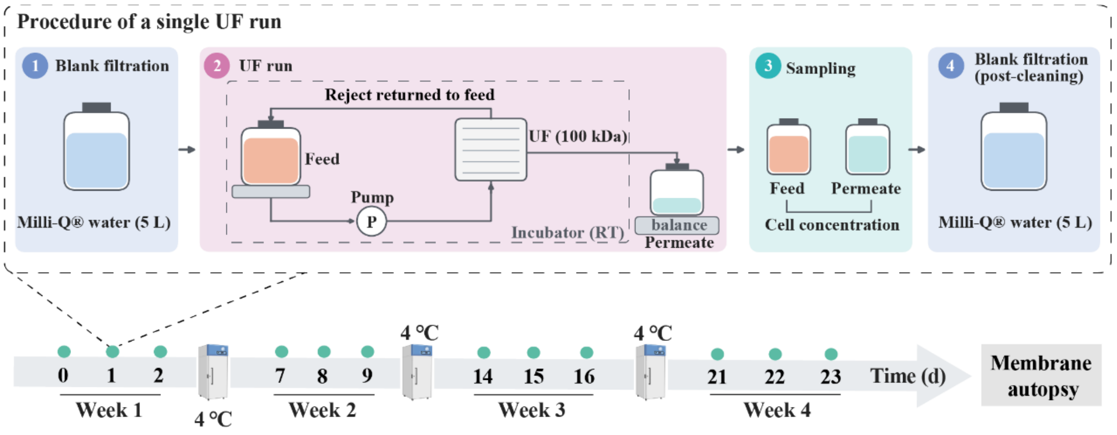


**Fig. S5** Experimental procedure of the pre-fouling experiment. Top: procedure of a single UF run, consisting of a blank filtration with 5 L of autoclaved Milli-Q® water, filtration of 4.3 L of fresh feed (*P. putida* K2440, $2.0 \times 10^7$ cells/mL in synthetic effluent), sampling for flow cytometry, and a second blank filtration. Bottom: timeline of the 12 UF runs over four weeks (not to scale). Between runs within a week, the cassette remained in the incubator at room temperature; after the third run of each week, the set-up was disassembled in a sterile bench, the membrane inlet and outlets were closed with sterile plugs, and the cassette was stored at 4 °C until the following week. Day 0: new membrane, no blank filtration.

### SI6. Flow cytometric measurement of cell numbers

Feed water and permeate samples were collected at the start (after 100 mL permeate had been collected) and at the end of the experiments to be analyzed with flow cytometry. The samples were preserved at 4 ºC with no fixation until SYTO®9 staining prior to flow cytometry performance. SYTO®9 (Lot. 2088729; ThermoFisher Scientific, Eugene, OR, USA) is a cell-permeant nucleic acid stain, which was used to distinguish fresh cells from environmental particles. A 35 µM stock solution of SYTO®9 was prepared daily with 0.85% NaCl (Sigma Aldrich). The final cell solution contained 950 µL cell sample and 50 µL 35 µM SYTO®9, with a final concentration of 1.75 µM SYTO®9. The cell solution was mixed and incubated for 15 min at RTin the dark before cell counting.

Cytometric measurement was performed with a BD Influx v7 Sorter USB, (Becton, Dickinson and Company Biosciences (BD), Franklin Lakes, USA) equipped with a blue 488 nm Sapphire OPSL (400 mW, Coherent, Santa Clara, USA). The 488 nm laser was used for excitation of forward scatter which is related to cell size (FSC (PMT1), band pass filter 488 nm ±5 nm), side scatter related to cell density (SSC (PMT2), trigger signal, band pass filter 488 nm ±5 nm), and the SYTO®9 fluorescence as nucleic acid stain (PMT3, band pass filter 530 nm ±20 nm). Light was detected by Hamamatsu R3896 photomultiplier (PMTs) in C6270 sockets (Hamamatsu, 211 Hamamatsu City, Japan). The fluidic system was run at 33 psi (2.275 bar) and a 70-µm nozzle. The sheath fluid consisted of FACSFlow solution (BD Biosciences). For calibration of the cytometric set up in the linear range, 1 µm blue fluorescent FluoSpheres F-8815 and 2 µm yellow-green fluorescent FluoSpheres F8827 (both ThermoFisher Scientific, Waltham, Massachusetts, USA) were used. For calibration of the logarithmic range 1 µm yellow-green fluorescent FluoSpheres F-13081 (Polysciences, Warrington, USA) were used. For cell analysis and counting, 1 µm yellow-green (YG) fluorescent FluoSpheres beads were added to every sample as control beads to ensure measurement stability and therefore allow comparison between samples. In addition they were used for cell counting as in Liu et al. (2019)[13].

The data were evaluated using FlowJo™ V10 software package (BD Biosciences), and a cell gate was generated following the general approach described by Cichocki et al. (2020)[14]. Beads and instrumental noise were excluded from the evaluation.

Control experiments were conducted to exclude events from counting that are no bacteria. The synthetic effluent medium can contain events such as salts[15-17] which could be detected by flow cytometry, causing a bias in the cell count measurements. Therefore, prior to conducting the experiments with bacteria, an analogous experiment was performed using only synthetic effluent medium to quantify the potential background coming from abiotic events. The background of the synthetic effluent medium was 4 orders of magnitude lower ($10^3$ events/mL) compared to the medium loaded with *P. putida* KT2440 and was therefore neglected (Fig. S6). After UF of the bacteria-free synthetic effluent medium, which was now called the permeate, showed the same low number of events (about $10^3$ events/mL) which was in this case subtracted as background signal from all measured permeates in the bacterial experiments.

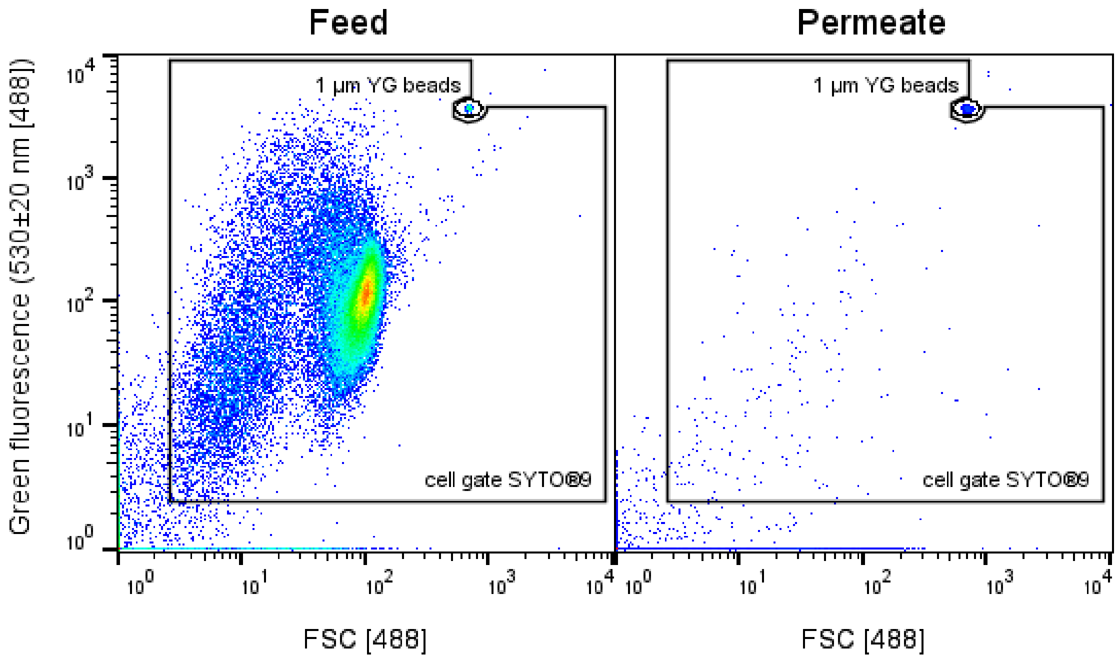


**Fig. S6** Flow cytometric controls for cell counting of *P.putida* KT2440 in synthetic effluent medium (feed) and permeate. Left: Setting of a cell gate for counting bacterial cells to exclude beads and instrumental noise. Right: Analysis of the synthetic effluent medium. For counting cells in the permeate, the number of events in the right cell gate were subtracted from the events in the left cell gate.

### SI7. Fouling visualization with HIM/SEM and Combined-FISH

The use of Helium Ion Microscopy (HIM) was planned to analyze the membrane surface after the experiments in Part I. However, after the first pre-fouling experiment (SI5), HIM was not available so Scanning Electron Microscopy (SEM) was used instead to verify the presence of bacteria in the membrane surface. Combined-FISH which integrates direct-gene Fluorescence in Situ Hybridization (FISH) for ARG detection with catalyzed reporter deposition fluorescence in situ hybridization (CARD-FISH) for rRNA labelling allowed the visualization of the plasmids deposited on the membrane surface.

Regardless of the microscopy technique applied, the autopsy procedure and sample treatment were the same: after the cassette opening under the sterile bench, the membrane sheets were cut in strips and immediately immersed in a 4% paraformaldehyde solution in PBS (Phosphate-buffered saline, 0.01 M, pH 7.2). The membrane fragments remained in the fixative solution overnight at 4ºC. Then, the fragments to be analyzed with HIM or SEM were subject to dehydration by consecutive immersion in the following solutions: 30% ethanol in PBS (5 min), 30% ethanol in Milli-Q® water (5 min), 50% ethanol in Milli-Q® water (5 min), 80% ethanol in Milli-Q® water (5 min), 90% ethanol in Milli-Q® water (5 min), 50% ethanol in hexamethyldisilazane (HMDS, 10 min) and 100% HMDS (10 min).

After the last step the membrane fragments were air dried and stored at 4ºC until further analyses. For Combined-FISH samples, the dehydration process was the same except for the fact that it stopped after immersion in 80% ethanol solution. After immersion in 80% ethanol solution, the membrane was air dried and small sections were then cut from the dehydrated membranes and used directly for Combined-FISH. Non-fouled membrane sections were only subjected to gene-targeted hybridization using the *sul1*-targeting polynucleotide probe set Sul1_3R. Fouled membrane sections were analyzed by Combined-FISH using Sul1_3R and the bacteria-targeted rRNA probe (EUB338I). Sul1_3R probes were directly labelled with Alexa Fluor 594 (Thermo Fisher Scientific, USA), whereas rRNA probe was labelled with horseradish peroxidase (HRP), and Alexa Fluor 488 was introduced by HRP-catalyzed deposition of Alexa Fluor 488-labelled tyramide. In addition, hybridization with the nonsense gene probe (Nonsul) and without any gene probe were used as negative controls. Sul1_3R and rRNA-targeted probe sequences, together with the hybridization procedures, were as described by in Wang et al. (2026) [18]. Prior to microscopy, hybridized membrane sections were secured on glass slides using nail polish. ProLong™ Gold Antifade Mountant (Thermo Fisher Scientific, USA) was then applied on the sample, and the sections were covered with coverslips, which were sealed in place with nail polish. Fluorescence microscopy was performed as described by Wang et al. (2026) [18].

### SI8. Additional calculations

1) Mass balance in the ultrafiltration membrane

The following equations can be used for the calculation of the mass balance[19] in the main experiments performed with plasmids as well as with the bacteria used for the fouling pre-experiment.

The feed water concentration at the start and at the end of the experiments, $C_{F0}$ and $C_{Fe}$, together with the permeate concentration, $C_p$, and the corresponding volumes $V_{F0}$, $V_{Fe}$ and $V_P$, allowed calculating the fractions of rejection and permeation with Eq. (S2) and (S3), respectively:

$$\text{Rejection [\%]} = \frac{C_{Fe} \cdot V_{Fe}}{C_{F0} \cdot V_{F0}} \cdot 100 \quad \text{(S2)}$$

$$\text{Permeation [\%]} = \frac{C_P \cdot V_P}{C_{F0} \cdot V_{F0}} \cdot 100 \quad \text{(S3)}$$

Being a cross-flow filtration system, Eq. (S4) would only be fulfilled if adsorption did not occur, as all the pollutants would either permeate or concentrate in the feed water.

$$C_{F0} \cdot V_{F0} = C_{Fe} \cdot V_{Fe} + C_p \cdot V_p \quad \text{(S4)}$$

On the contrary, if a certain percentage was adsorbed on the membrane surface, it could be calculated in accordance with Eq. (S5).

$$\text{Adsorption [\%]} = \frac{C_{F0} \cdot V_{F0} - C_{Fe} \cdot V_{Fe} - C_p \cdot V_p}{C_{F0} \cdot V_{F0}} \cdot 100 \quad \text{(S5)}$$

The total removal can be calculated as the sum of the percentage of rejection and the percentage of adsorption and should be coincident with that in Eq. (S6).

$$\text{Total removal [\%]} = \left(1 - \frac{C_P \cdot V_P}{C_{F0} \cdot V_{F0}}\right) \cdot 100 \quad \text{(S6)}$$

2) Fouling calculation in terms of membrane resistance

Fouling can be calculated indirectly in terms of membrane resistance with the variation of transmembrane pressure (TMP) or flux ($J$) throughout the experiments.

This method is approximate and has some limitations, e.g., it depends on the intensity of the cleaning procedure. Furthermore, it can only be used when the permeate flow rate (at constant TMP) or TMP (at constant flux) changes significantly over the course of the experiments, so it is not useful when working with very diluted feed water. However, for filtration of bacterial solutions with a fixed cleaning routine, as in the case of the pre-fouling experiments, it can give an idea of how quickly the membrane becomes fouled in the course of the experiments.

The permeate flow-rate was registered by an electronic balance connected to a computer throughout the experiments, for which the flux, $J$ ($m^3/m^2 \cdot s$) was calculated as the ratio between the permeate flow rate ($Q_p$, $m^3/s$) and the membrane area ($A_m$, $m^2$), Eq. (S7).

$$J = \frac{Q_p}{A_m} \quad \text{(S7)}$$

The transmembrane pressure, TMP (Pa), for a crossflow filtration system is given by Eq. (S8), with feed and reject pressures, $P_{feed}$ and $P_{reject}$, being measured by respective pressure gauges. Permeate pressure, $P_p$ is null in our system.

$$TMP = \frac{P_{feed}+P_{reject}}{2} - P_p \quad \text{(S8)}$$

The membrane resistance, $R$ ($m^{-1}$) can be calculated for the corresponding flux and TMP conditions together with the dynamic viscosity of the permeate, $\mu$ (Pa·s), with Eq. (S9):

$$R = \frac{TMP}{\mu \cdot J} \quad \text{(S9)}$$

Additionally, the total membrane resistance has several components, as indicated in Eq. (S10):

$$R_t = Rm + Ri + Rr \quad \text{(S10)}$$

$R_m$ is the intrinsic membrane resistance, calculated after membrane compaction with autoclaved Milli-Q® water. $R_i$ and $R_r$ represent, respectively, the irreversible and reversible components of the membrane resistance[20-22].

As indicated in the Materials and Methods section, a blank filtration with 5L autoclaved Milli-Q® water was performed before and after each ultrafiltration run with feed water (either with bacteria or plasmids experiments) to flush the membrane surface and avoid bias between experiments.

If all the reversible fouling is assumed to be removed with the membrane cleaning[20-21], the residual total resistance after cleaning, $R_t^{'}$, is only composed of $R_i$ and $R_m$ (Eq. (S11)).

$$R_t^{'} = Rm + Ri \tag{S11}$$

Since $R_m$ is known, the calculation of $R_t^{'}$ with Eq. S10 for the conditions after cleaning allows to obtain $R_i$. In turn, the calculation of $R_t$ with Eq. S9 at the end of each filtration experiment allows the calculation of $R_r$. The evolution of $R_i$ and $R_r$ over the experiments can give an idea of what is happening in the membrane without the need to open the membrane cassette, which would be irreversible for this type of assembly.
